# 14-helical β-peptides Elicit Toxicity against *C. albicans* by Forming Pores in the Cell Membrane and Subsequently Disrupting Intracellular Organelles

**DOI:** 10.1101/430850

**Authors:** Myung-Ryul Lee, Namrata Raman, Patricia Ortiz-Bermudez, David M. Lynn, Sean P. Palecek

**Author notes:** To whom correspondence should be addressed: David M. Lynn, 1008 Engineering Hall, 1415 Engineering Drive, Madison, WI 53706-1691, Sean P. Palecek (Lead Contact), 3637 Engineering Hall, 1415 Engineering Drive, Madison, WI 53706-1691.

## Abstract

Synthetic peptidomimetics of antimicrobial peptides are promising as antimicrobial drug candidates because they promote membrane disruption and exhibit greater structural and proteolytic stability. We previously reported selective antifungal 14-helical β-peptides, but the mechanism of antifungal toxicity of β-peptides remains unknown. To provide insight into the mechanism, we studied antifungal β-peptide binding to artificial membranes and living *Candida albicans* cells. We investigated the ability of β-peptides to interact with and permeate small unilamellar vesicle models of fungal and bacterial membranes. The partition coefficient supported a pore-mediated mechanism characterized by the existence of a critical β-peptide concentration separating low and high partition coefficient regimes. Live cell intracellular tracking of β-peptides showed that β-peptides translocated into the cytoplasm, and then disrupted the nucleus and vacuole sequentially, leading to cell death. This understanding of the mechanisms of antifungal activity will facilitate design and development of peptidomimetic AMPs, including 14-helical β-peptides, for antifungal applications.

**Graphical Abstract:** 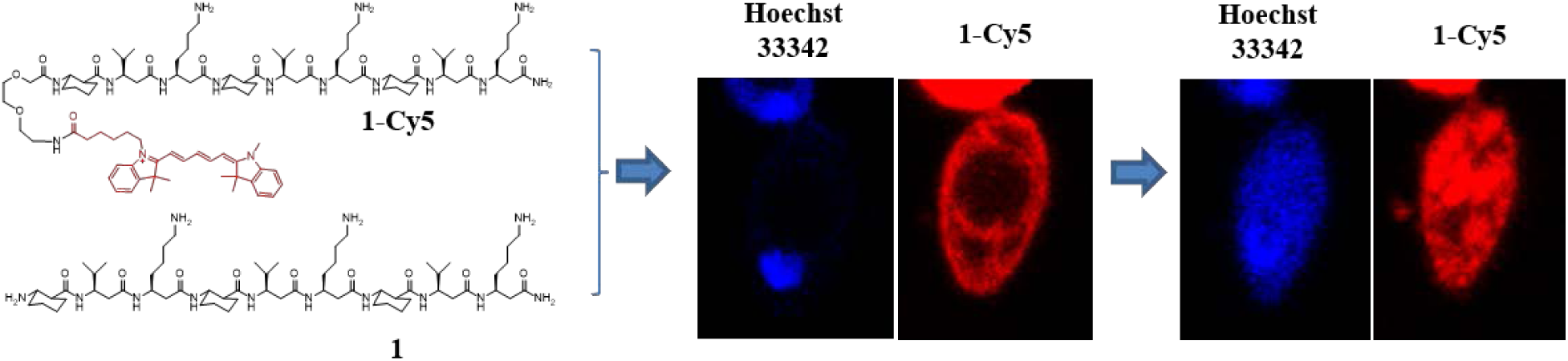

## Introduction

Antimicrobial peptides (AMPs) are components of innate immune systems found in a wide variety of invertebrates and vertebrates (Beutler, 2004; Gallo and Nizet, 2003; Ganz, 2003), and have demonstrated broad spectrum activity against bacteria, fungi, viruses, and cancer cells (Brogden et al., 2003; Hancock and Lehrer, 1998; Zasloff, 2002). Only a few naturally-occurring instances of resistance to AMPs have been reported (Andersson and Hughes, 2010; Nizet, 2006). Thus, there is significant interest in developing AMPs as drugs against infectious agents since many existing antimicrobials are plagued by the emergence of resistant strains. Unfortunately, the viability of natural AMPs as therapeutics is reduced by their low structural stability and activity in physiologic environments (Chu et al., 2013; Goldman et al., 1997; Huang et al., 2011; Smith et al., 1996) and their susceptibility to proteolytic cleavage (Schmidtchen et al., 2002).

A variety of synthetic or non-natural peptidomimetic compounds have been developed to exploit the antimicrobial properties of AMPs and to address their low stability (Godballe et al., 2011; Lienkamp et al., 2013). One such class of compounds, β-peptides, is composed of β-amino acids and can fold into protein secondary structures found in AMPs, including the α-helix (Appella et al., 1997; Seebach et al., 1997; Seebach and Matthews, 1997), β-sheet (Chung et al., 2000; Krauthauser et al., 1997), and γ-turn (Chung et al., 1998). Structure-functional analyses of β-peptides have demonstrated that these compounds can be designed to exhibit antibacterial (Arvidsson et al., 2001; Epand et al., 2003; Hamuro et al., 1999; Porter et al., 2000; Raguse et al., 2002) and antifungal (Karlsson et al., 2009; Raman et al., 2015) activities at concentrations that induce little hemolysis in mammalian red blood cells. In prior work, we demonstrated that fungicidal activity of 14-helical β-peptides against the opportunistic pathogen *Candida albicans* requires a globally-amphiphilic structure, composed of hydrophobic and cationic faces, with approximately three helical turns (Lee et al., 2014). We observed that both antifungal and hemolytic activities increased as β-peptides became more hydrophobic, and we identified a hydrophobicity window that resulted in high antifungal activity but low hemolysis. The addition of helix-stabilizing residues, such as aminocyclohexane carboxylic acid (ACHC), increased the activity and specificity of these 14-helical β-peptides against *C. albicans*.

While prior studies have shown that β-peptides and other AMP peptidomimetics have promise as antimicrobial therapeutics, little is known about their mechanism of action. The canonical mechanism of AMP-mediated toxicity involves permeabilization of the target cell membrane, a process that depends on both the physicochemical properties of the AMP and those of the target cell membrane. Several models have been proposed to describe the mechanisms of AMP permeabilization of cell membranes (Brogden, 2005; Huang, 2000; Shai, 1999; Yeaman and Yount, 2003). In the “barrel-stave model”, AMP monomers bind to the cell membrane and then oligomerize and form structured pores with the hydrophobic region of the peptide oriented toward the membrane lipids and the hydrophilic region comprising the pore channel. For example, aggregation of four to six alamethicin molecules in a cell membrane has been shown to form voltage-dependent ion channels (Boheim, 1974; Gordon and Haydon, 1972, 1975). In contrast, in the “toroidal model”, peptides insert into the cell membrane and associate with lipid headgroups to induce lipid bending. Thus, the pore lining is composed of both the peptides and membrane lipids. Magainin 2 (Matsuzaki et al., 1995; Yang et al., 1998), protegrin-1 (Lazaridis et al., 2013; Yamaguchi et al., 2002), melittin (Yang et al., 2001; Zhou et al., 2014), LL-37 (Lee et al., 2011), MSI-78 (Hallock et al., 2003), and other AMPs have been shown to act via the toroidal model. Finally, in the “carpet model”, AMPs first interact with the target membrane via electrostatic interactions, and then they effectively solubilize and disrupt the membrane. Dermaceptin S (Pouny et al., 1992), cecropin (Gazit et al., 1996), caerin 1.1 (Pukala et al., 2004), ovispirin (Yamaguchi et al., 2001), and other AMPs have been suggested to follow the carpet model. Little is known about how antifungal peptidomimetic 14-helical β-peptides may induce membrane lysis and promote cell death in *C. albicans*.

In addition to permeabilizing the cell membrane, some AMPs act on intracellular targets to induce microbial death. Modes of intracellular action include binding nucleic acids (e.g. buforin II (Park et al., 1998) and tachyplesin (Yonezawa et al., 1992)) and inhibiting cell-wall synthesis (mersacidin (Brotz et al., 1995)), nucleic acid and protein synthesis (pleurocidin (Patrzykat et al., 2002), PR-39 (Boman et al., 1993), and HNP-1 (Lehrer et al., 1989)), and enzymatic activity (histatins (Nishikata et al., 1991; Puri and Edgerton, 2014), pyrrhocoricin (Gagnon et al., 2016; Otvos et al., 2000), and drosocin (Otvos et al., 2000)).

Prior studies have focused on interaction of AMPs with the cell membrane, but their effects on intracellular compartments, such as the nucleus, vacuole, and mitochondria have not been studied. In addition, there is little understanding of membrane disrupting mechanism of peptidomimetic β-peptides and the fates of these compounds inside the target cell. Here, we studied interactions between 14-helical β-peptides and small unilamellar vesicles (SUVs) to elicit the membrane disrupting mechanism and tracked fluorescently-labeled β-peptide to understand the interactions between peptide and intracellular organelles.

To elucidate mechanisms of β-peptide interactions with fungal cells, we first employed SUVs to investigate how membrane composition influences β-peptide adhesion to and the subsequent permeabilization of the vesicles. Next, we used confocal laser scanning microscopy (CLSM) and super resolution structure illumination microscopy (SR-SIM) to validate that cooperative binding of β-peptides observed in SUVs also occurs in *C. albicans* cells. Finally, time lapse microscopy tracking of fluorescently labelled β-peptides demonstrated that, after permeabilizing the cell membranes, the β-peptides lysed the nucleus and vacuole, resulting in cell death. Our results are consistent with pore-mediated mechanisms of membrane permeabilization and suggest that disruption of intracellular organelles also plays a role in the fungicidal activity of β-peptides. The improved understanding of mechanisms of β-peptide interactions with the cell membrane and organelles will enhance efforts to further develop these compounds as active and selective antifungal agents.

## RESULTS

### Hydrophobicity of β-peptide regulates the antifungal activity

To study the potential mechanisms of antifungal β-peptide toxicity, we synthesized β-peptides **1** (NH_2_-(ACHC-β^3^hVal-β^3^hLys)_3_-CONH_2_), **2** (NH_2_-(ACHC-β^3^hVal-β^3^hArg)_3_-CONH_2_) and **3** (NH_2_-(ACHC-β^3^hAal-β^3^hLys)_3_-CONH_2_) (Fig. 1). We previously demonstrated that β-peptides **1, 2**, and **3** share structural properties important in antifungal activity, including charge, number of β-amino acid residues, amphiphilicity, and helical stability (Lee et al., 2014; Raman et al., 2015). However, the hydrophobicity of these β-peptides (as determined by RP-HPLC retention time) (Browne et al., 1982; Jiang et al., 2008; Kondejewski et al., 1999; Wilson et al., 1981), which also regulates the antifungal activity, varies substantially. The more hydrophobic β-peptides **1** and **2** exhibit an MIC against *C. albicans* of 8 μg/mL, while the MIC of less hydrophobic β-peptide **3** is 512 μg/mL (Table 1). We also synthesized NBD-labeled analogues of β-peptides **1, 2**, and **3** (**1-NBD**, **2-NBD**, and **3-NBD**) and Cy5-labeled β-peptide **1** (**1-Cy5**) (Fig. 1) to facilitate quantification of β-peptide partition coefficients into synthetic lipid vesicle membranes and to visualize the interactions between the β-peptides and live *C. albicans* cells, respectively. After NBD-labeling of β-peptides **1**, **2**, and **3**, peptide retention times increased, indicating that the NBD label increased the hydrophobicity of these β-peptides (Table 1, Fig. S1). The MICs of β-peptides **1-NBD** and **2-NBD** were within a factor of 2 of the MICs of the corresponding unlabeled β-peptides (Table 1, Fig. S2) while the MIC of **3-NBD** was 8-fold lower than the MIC of β-peptide **3** (Table 1, Fig. S2), likely resulting from the increase in β-peptide hydrophobicity upon adding the NBD label. β-peptide **3-NBD** still exhibited a 4- and 8-fold greater MIC than **1-NBD** and **2-NBD**, respectively. To characterize mechanisms of antifungal β-peptide toxicity, we investigated biophysical interactions of fluorescently-labelled β-peptides with model phospholipid membranes and *C. albicans* cell membranes, as well as the subsequent localization of β-peptides to intracellular compartments.

**Figure 1.**
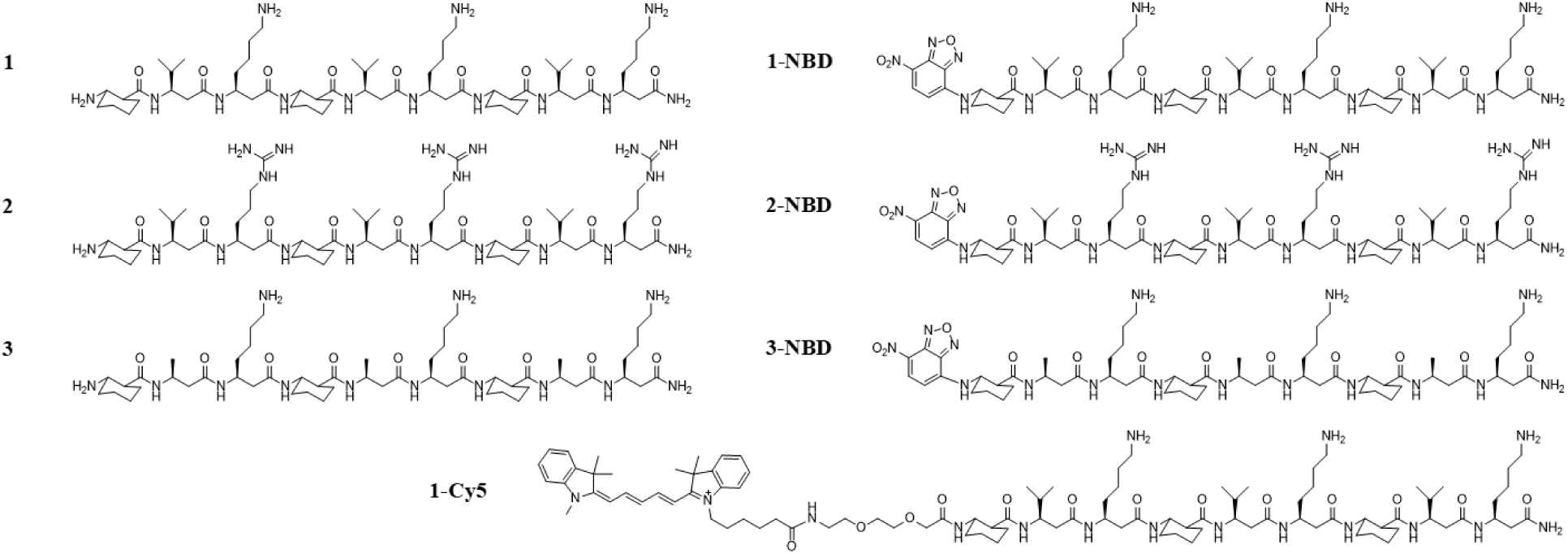
Structures of β-peptides used in this study.

**Table 1.** Retention times and minimum inhibitory concentrations (MIC) against *C. albicans* for β- peptides used in this study.

| Peptide | RT<sup>a</sup><br>(min ± SD) | MIC<sup>b</sup><br>(μg/mL) | MIC<sup>b</sup><br>(μM) | Peptide | RT<sup>a</sup><br>(min ± SD) | MIC<sup>b</sup><br>(μg/mL) | MIC<sup>b</sup><br>(μM) |
| --- | --- | --- | --- | --- | --- | --- | --- |
| 1 | 21.13 ± 0.04 | 8 | 5.0 | 1-NBD | 29.43 ± 0.08 | 16 | 9.6 |
| 2 | 22.03 ± 0.07 | 8 | 4.7 | 2-NBD | 31.04 ± 0.07 | 8 | 4.6 |
| 3 | 15.49 ± 0.01 | 512 | 334.5 | 3-NBD | 21.07 ± 0.04 | 64 | 40.5 |
| Magainin 2 | | 512 | 3390.2 | 1-Cy5 | 28.89 ± 0.05 | 4 | 1.9 |
| Melittin |  | 4 | 1.1 |  |  |  |  |

<sup>a</sup> The average value obtained from three independent analytical RP-HPLC measurements. <sup>b</sup> The median value obtained from three independent experiments with triplicate measurements in each. SD denotes standard deviation (n = 3).

### Permeabilization of synthetic vesicles by β-peptides depends upon membrane composition

We used the leakage of calcein from synthetic small unilamellar vesicles (SUVs) to assess the specificity of β-peptide disruption of phospholipid membranes composed of phosphatidylglycerol (PG, anionic phospholipid), phosphatidylcholine (PC, neutral phospholipid), and phosphatidylethanolamine (PE, neutral phospholipid). We prepared SUVs having different ratios of these phospholipids including PC:PE (60:40), PC:PE (80:20) as a model for fungal cell membranes (Warschawski et al., 2011), and PC:PG (60:40) as a model for a bacterial membrane (Tamba and Yamazaki, 2005). After preparation, the SUV size and phospholipid ratio were quantified using dynamic light scattering and ^31^P nuclear magnetic resonance (Table S1). Magainin 2, a pore-forming antibacterial peptide, and melittin, a pore-forming non-selective AMP, were used as controls for membrane lysis. Magainin 2 induced substantial calcein leakage only from the model bacterial membrane vesicles (PC:PG 60:40) (Fig. 2, and S3-5). In contrast, melittin induced calcein leakage from all SUVs tested (Fig. 2, and S3-5), as expected given its broad activity against bacterial, fungal, and mammalian cells. The phospholipid selectivities of magainin 2 and melittin for SUVs are consistent with prior reports of PG selectivity of magainin 2 (Gregory et al., 2009; Wieprecht et al., 1996) and broad phospholipid targeting of melittin (Magzoub et al., 2005) against large unilamellar vesicles (LUVs).

**Figure 2.**
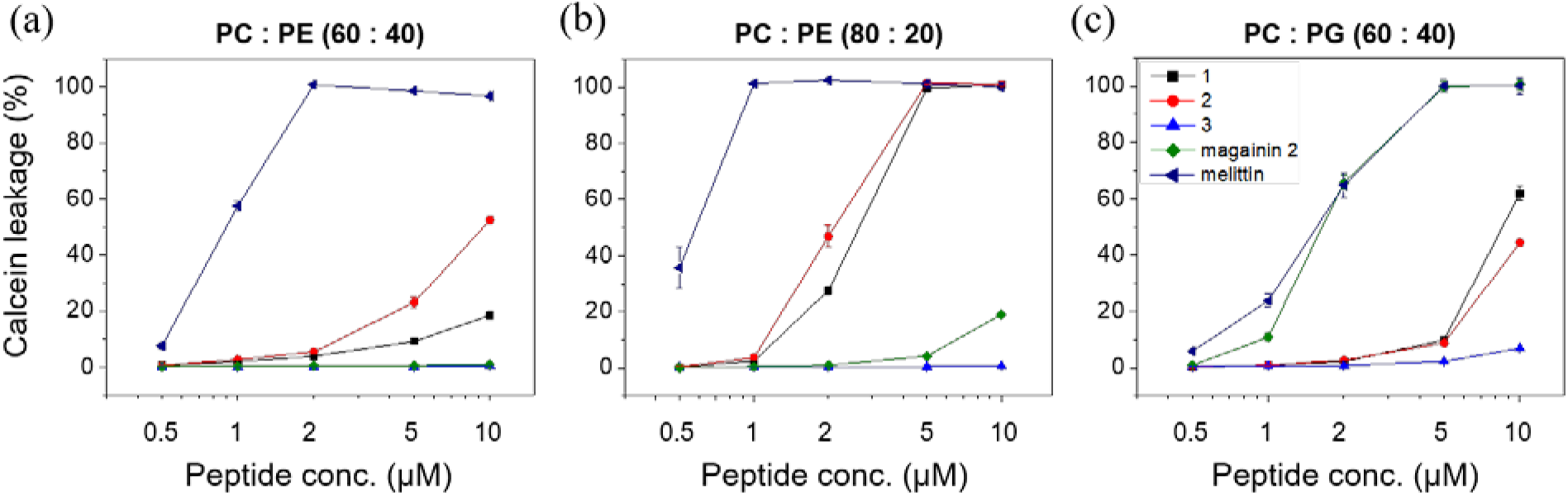
β-Peptide concentration-dependent calcein leakage from SUVs of the indicated phospholipid concentrations. Calcein leakage was monitored 10 min after incubation of β-peptides and SUVs (total [lipid] = 130 μM). To quantify calcein leakage, the fluorescence intensity at 515 nm was measured and normalized to the intensity obtained after adding 2% Triton X-100, corresponding to 100% calcein leakage. Error bars denote standard deviation (n = 3).

The disruption of SUVs by 14-helical β-peptides depended on the SUV and β-peptide compositions. The active antifungal β-peptides **1** and **2** resulted in about 10–20% calcein leakage from PC:PE (60:40) SUVs and 100% calcein leakage from PC:PE (80:20) SUVs after 10 min exposure to the β-peptides at the antifungal MIC (5 μM, Table 1) (Fig. 2, Fig. S3 - 4). The more hydrophobic β-peptide **2** induced slightly more calcein leakage from the PC:PE (60:40) SUVs than did β-peptide **1**. In contrast, the inactive β-peptide **3** resulted in less than 2% calcein leakage even after 1 hr from neutral SUVs (Fig. 2, Fig. S3 - 4).

We also quantified calcein leakage from negatively-charged PC:PG (60:40) SUVs to investigate the effects of electrostatic interactions on phospholipid membrane permeabilization by 14-helical β-peptides. β-peptides **1** and **2** lysed PC:PG (60:40) SUVs to a substantially greater extent than β-peptide **3** at and above 5 μM (Fig. 2c). However, β-peptides **1** and **2** elicited similar calcein leakage in PC:PG (60:40) SUVs as PC:PE (60:40) SUVs at 10 μM. β-peptide **3** caused more than ten-fold greater calcein leakage after 1 h from PC:PG (60:40) SUVs than from PC:PE (60:40) SUVs (Fig. S5). The activity of hydrophobic β-peptides **1** and **2** appeared to be less sensitive to electrostatic interactions than the less hydrophobic β-peptide **3**. Together, these results demonstrate that the ability of 14-helical β-peptides to lyse model microbial cell membranes correlates with their antifungal activity, and suggest a potential mechanism of antifungal activity that involves cell membrane disruption.

### 14-Helical β-peptides lyse SUV membranes via pore formation

To characterize mechanisms of membrane disruption and distinguish between pore-forming and carpet mechanisms, we quantified the partition coefficients of NBD-labeled β-peptides in model SUV membranes. Carpet formation is characterized by a monotonic correlation between peptide concentration and membrane binding (Rapaport and Shai, 1991). In contrast, a hallmark of pore formation is two regimes of peptide binding to the membrane (Huang, 2000; Jackman et al., 2016). Below the critical concentration, a low partition coefficient results from peptide interactions with the membrane surface, while above the critical concentration a higher partition coefficient results from cooperative insertion of the peptides into the membrane as pores are formed (Brogden, 2005; Huang, 2000; Shai, 1999; Yeaman and Yount, 2003).

We measured partition coefficients of **1-NBD**, **2-NBD** and **3-NBD** in PC:PE (60:40), PC:PG (60:40), and PC:PE:ergosterol (50:30:20) SUVs (Table 2, and S2, and Fig. 3, and S6 - 9). Partition coefficients were obtained by plotting *X*_b_^*^, the corrected molar ratio of bound peptide per total outer leaflet lipid (assumed to be 60% of the total lipid) (Michaelson et al., 1974) versus *C*_f_, the equilibrium concentration of free peptide in solution. The partition coefficient was calculated from the slope of this curve (Fig. 3, and S7 - 9). **1-NBD** and **2-NBD**, which exhibit high antifungal activity (Table 1), demonstrated greater partition coefficients, 23.6- and 35-fold higher respectively, than the less active and more hydrophilic **3-NBD** in PC:PE (60:40) SUVs (Table 2, Fig. 3). The partition coefficients of **1-NBD** and **2-NBD** varied by 2-fold or less between PC:PG SUVs, while **3-NBD** exhibited a 16-fold greater partition coefficient in PC:PG SUVs than PC:PE SUVs, indicating a greater dependence of partition coefficient on electrostatic interactions for **3-NBD** (Table 2, Fig. 3). As anticipated, the partition coefficients for NBD-labeled β-peptides correlated with the extent of calcein leakage caused by the unlabeled β-peptides (Fig. 2). Critical concentrations of 13.7 and 17.8 nM were identified for **1-NBD** and **2-NBD**, respectively, for PC:PE (60:40) SUVs, but **3-NBD** did not exhibit a critical concentration against PC:PE SUVs, consistent with a mechanism of pore formation by **1-NBD** and **2-NBD** but not **3-NBD**. All three NBD-labeled β-peptides exhibited critical concentrations between 11 and 16 nM against PC:PG (60:40) SUVs, suggesting that all three NBD-labeled β-peptides formed pores in PC:PG (60:40) SUVs (Table 2).

**Figure 3.**
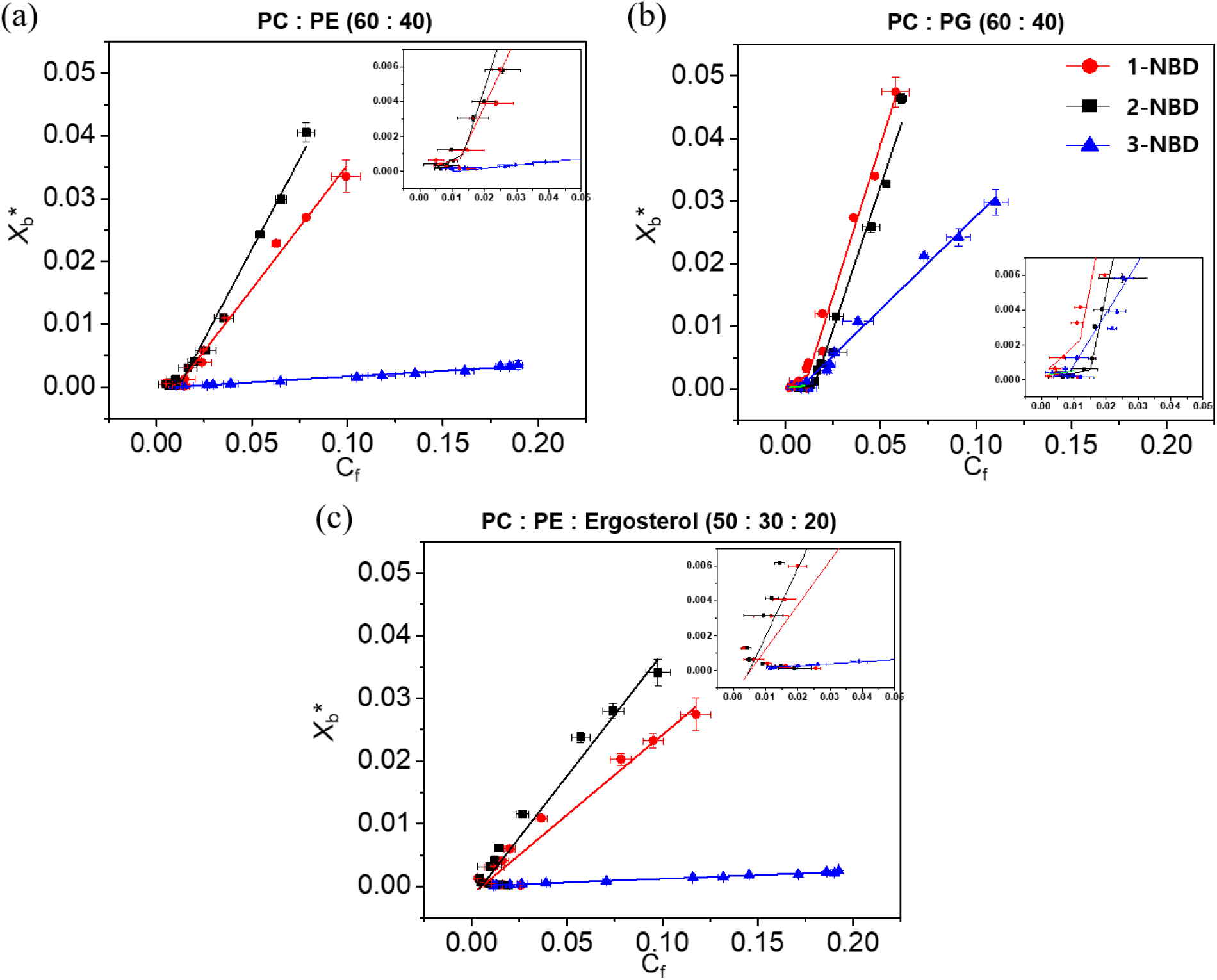
Binding isotherms of NBD-labeled β-peptides (0.2 μM) to various concentrations of SUVs ([lipid] = 5 to 2000 μM) with the indicated compositions after 2 min incubation. *X*_b_^*^ represents the molar ratio of bound peptide per 60% of total lipid and *C_f_* is the equilibrium concentration of free peptide in solution. The partition coefficient is the slope of the binding isotherm and the critical concentration is the intersection of the low-slope and high-slope isotherms, shown in the insets. Data points indicate the averages of three independent experiments and error bars denote standard deviation (n = 3).

**Table 2.**
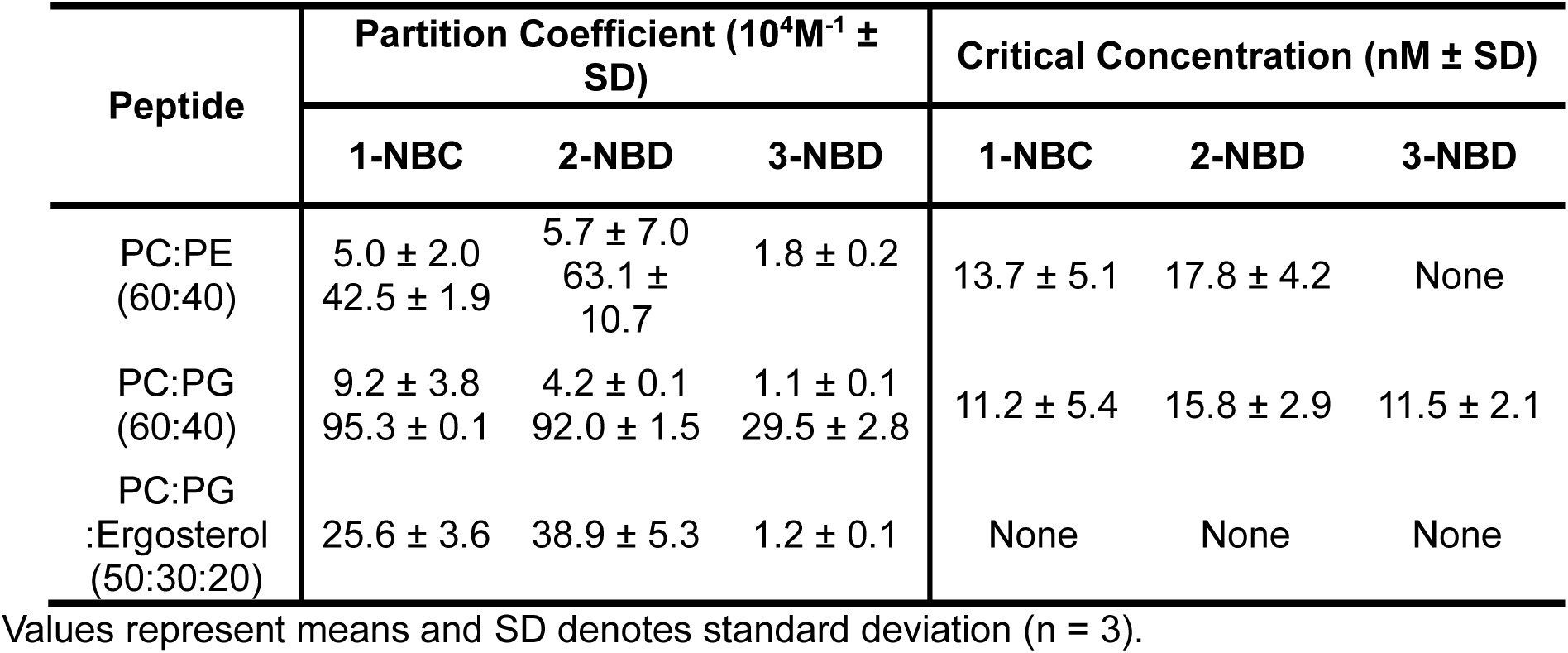
Partition coefficients and critical concentrations obtained for NBD-labeled β-peptides with SUVs of the indicated lipid compositions.

| Peptide | Partition Coefficient ($10^4 M^{-1} \pm$ SD) | | | Critical Concentration (nM ± SD) | | |
| --- | --- | --- | --- | --- | --- | --- |
|  | 1-NBC | 2-NBD | 3-NBD | 1-NBC | 2-NBD | 3-NBD |
| PC:PE (60:40) | 5.0 ± 2.0<br>42.5 ± 1.9 | 5.7 ± 7.0<br>63.1 ± 10.7 | 1.8 ± 0.2 | 13.7 ± 5.1 | 17.8 ± 4.2 | None |
| PC:PG (60:40) | 9.2 ± 3.8<br>95.3 ± 0.1 | 4.2 ± 0.1<br>92.0 ± 1.5 | 1.1 ± 0.1<br>29.5 ± 2.8 | 11.2 ± 5.4 | 15.8 ± 2.9 | 11.5 ± 2.1 |
| PC:PG :Ergosterol (50:30:20) | 25.6 ± 3.6 | 38.9 ± 5.3 | 1.2 ± 0.1 | None | None | None |

Values represent means and SD denotes standard deviation (n = 3).

We also investigated the effects of ergosterol on β-peptide partition coefficients in PC: PE SUVs (Fig. 3c). The ratio of ergosterol was adjusted to mimic that found in the *C. albicans* cell membrane (Warschawski et al., 2011). The partition coefficients of all NBD-labeled β-peptides in PC:PE:ergosterol (50:30:20) were lower than in PC: PE (60:40) SUVs (Fig. 3c, Table 2). The membrane sterols may inhibit the β-peptide adhesion to the membrane, as previously suggested (Bennett et al., 2009; Mason et al., 2007). These results indicate that membrane composition dictates the ability of NBD-labeled β-peptides to form pores in these model SUVs.

### 14-Helical β-peptides are cooperatively inserted into the plasma membrane of *C. albicans*

The calcein leakage and partition coefficient experiments suggest that antifungal β-peptides permeate synthetic phospholipid membranes by binding to and forming pores in the membranes. To explore mechanisms of β-peptide permeation of *C. albicans* cell membranes, we used confocal laser scanning microscopy (CLSM) and image analysis to measure relative binding of β-peptides to live *C. albicans* cells as a function of β-peptide concentration. In these experiments, the concentration of fluorescently-labeled β-peptide **1-Cy5** was fixed at 0.5 μM and unlabeled β- peptide **1** was added to vary the total β-peptide concentration from 0.5 to 7.0 μM. The cell-associated fluorescence signal was significantly greater above the MIC of β-peptide **1** (5.0 μM) than below the MIC (Fig. 4 and Fig. S10), suggesting cooperative binding of the β-peptide to the *C. albicans* cell membrane. These results are consistent with a pore-forming mechanism of fungal cell toxicity and suggest a similar mechanism of poration of SUVs and living *C. albicans* cell membranes.

**Figure 4.**
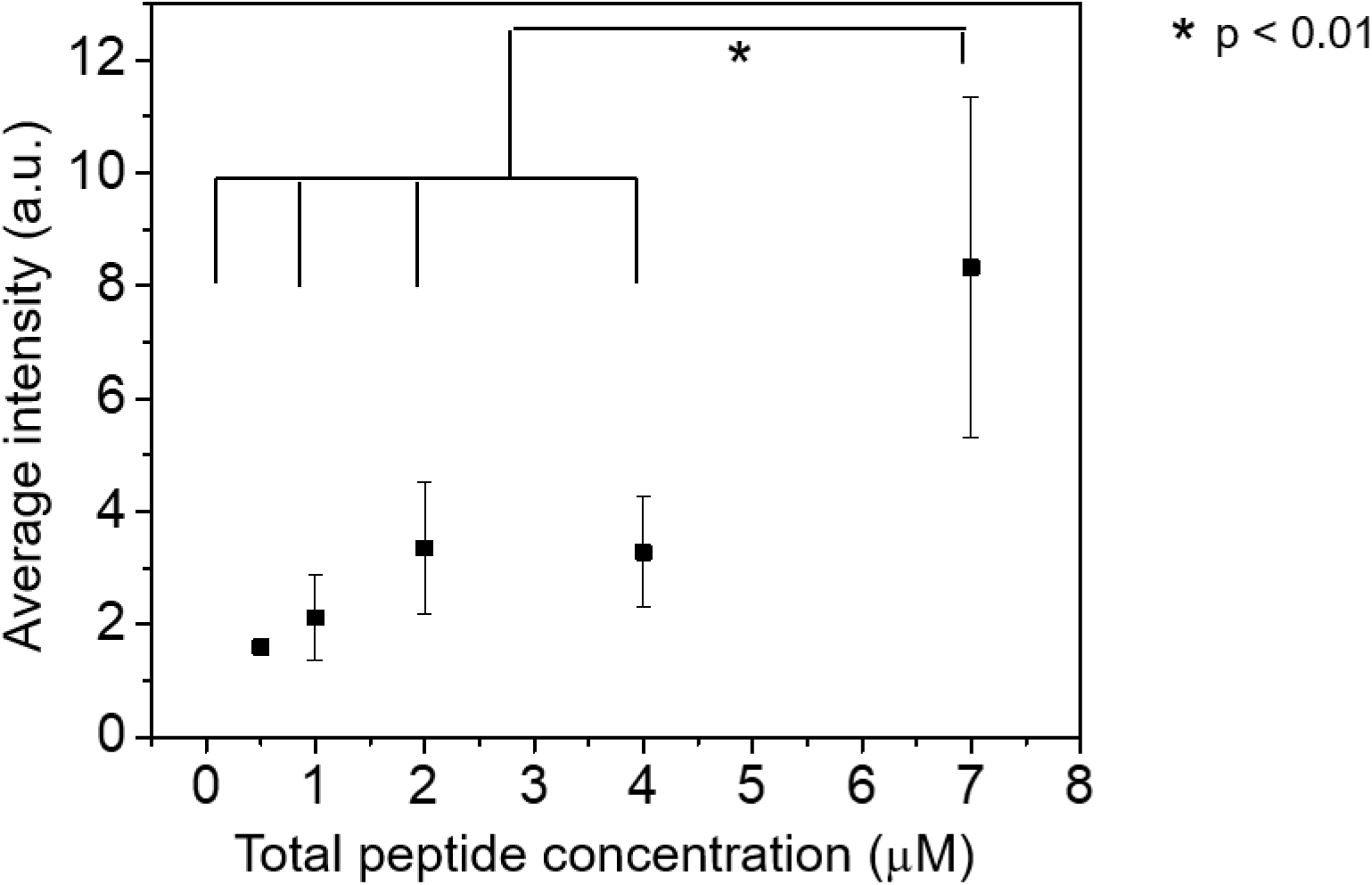
Cooperative insertion of β-peptide 1-Cy5 in the C. *albicans* plasma membrane. Cy5 fluorescence intensity associated with each cell was measured by fluorescence microscopy after 3 min incubation with a solution of 0.5 μM β-peptide 1-Cy5 and 0–6.5 μM β-peptide 1 in DPBS. The fluorescence intensity was measured for individual cells and then background intensity subtracted (Fig. S10). Data points are the averages of three independent experiments with approximately 20 cells/experiment, and error bars denote standard deviation (n = 3). Statistical analysis was performed using one-way ANOVA.

We next used super-resolution structure illumination microscopy (SR-SIM) to visualize β- peptide distribution on the *C. albicans* membranes. A sub-MIC concentration of β-peptide **1-Cy5** (2 μM) was incubated with live *C. albicans* for 3 min then imaged. Fig. 5 demonstrates that the β- peptides congregated into punctate structures around 300 nm diameter on the cell surface (Fig. 5). This result suggests rapid adhesion and aggregation of β-peptides on the plasma membrane. The concentrated β-peptides in these structures likely facilitated the supramolecular aggregation between β-peptide and lipids on the plasma membrane surface, which enables cooperative insertion into the plasma membrane to form pores in the *C. albicans* cell membrane.

**Figure 5.**
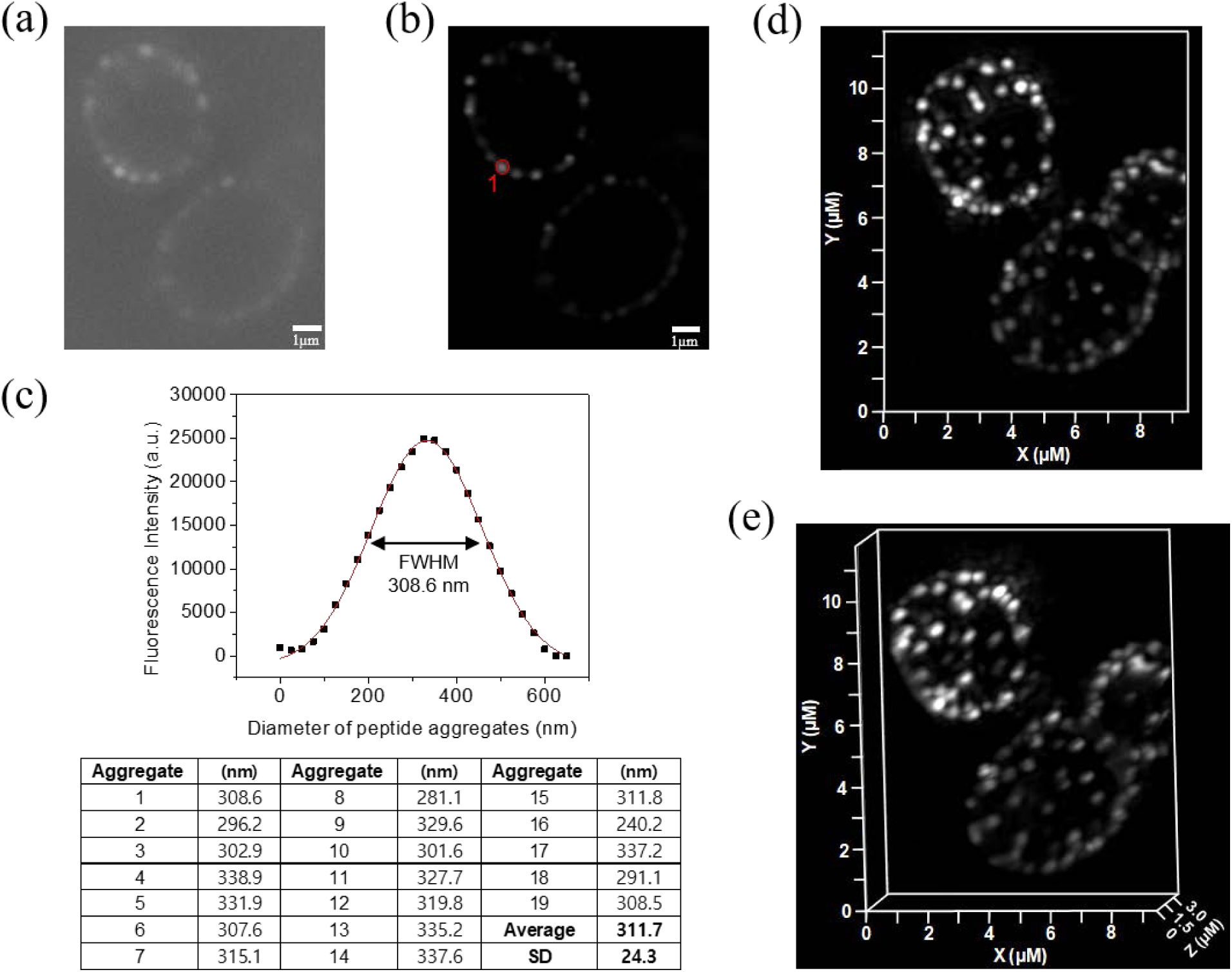
SR-SIM of β-peptide aggregation on the plasma membrane of *C. albicans*. 2 μM β- peptide 1-Cy5 was added to *C. albicans* cells and cells were imaged after 3 min incubation. Panel (a) shows a sample raw z-section image while panel (b) shows the same image processed using Zeiss ZEN software. (c) Fluorescence intensity distribution of the peptide aggregate circled in red in (b). To measure the size of peptide aggregate, the fluorescence intensity vs diameter plot was fit to a Gaussian distribution (red line). The full width at half maximum intensity was taken to the be aggregate size. Average peptide aggregate size was calculated from 19 aggregates from 5 different cells. SD denotes standard deviation (n = 19). Panels (d) and (e) show 3D-SR SIM image projections of stacked of Z-sections. Scale bar indicates 1 μm for (a) and (b).

### 14-Helical β-peptides form PI-permeable pores on live *C. albicans*

Figures 3–5 suggest that β-peptides form pores in model phospholipid membranes and in living *C. albicans* cell membranes. Next, we investigated whether these pores facilitate molecular translocation across the cell membrane and into the cytoplasm using the membrane-impermeable DNA-binding fluorescent dye propidium iodide (PI) as a tracer. We incubated *C. albicans* cells in sub-MIC (0–3 μM) concentrations of β-peptide **1** in the presence of 10 μg/mL PI and monitored PI accumulation inside the cell for 45 min using CLSM. At 2–3 μM β-peptide **1**, some cells were lysed, indicated by high intensity PI staining (red cells in Fig. 6a and S11). Quantification of PI staining in the living cells demonstrated that as β-peptide concentration increased, the PI uptake into living cells increased throughout the incubation period (Fig. 6b). These results suggest that β-peptides can form pores in the plasma membrane that permit molecular translocation at concentrations below the MIC.

**Figure 6.**
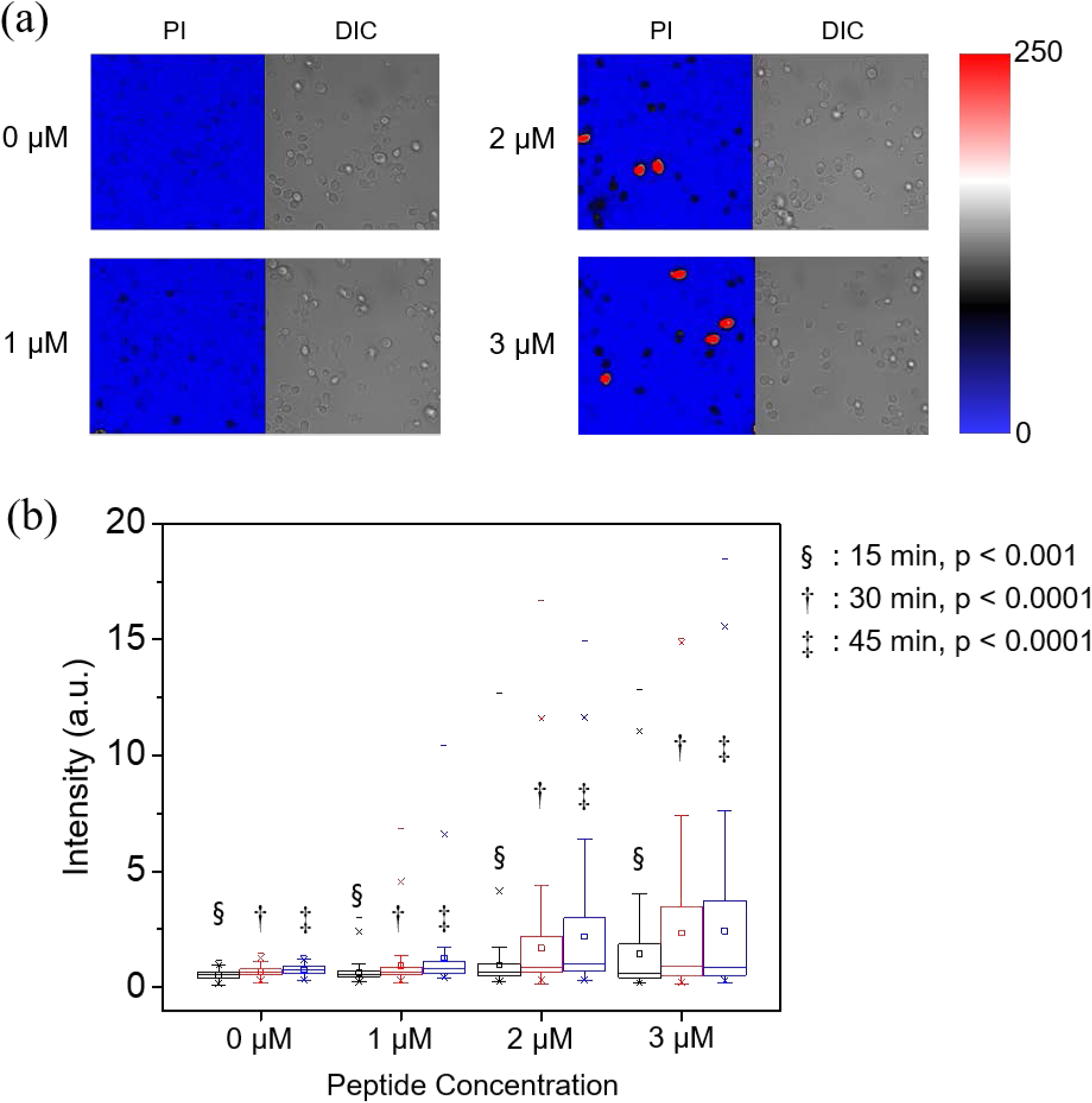
PI uptake into live *C. albicans*. Cells were incubated with 10 μg/mL PI and the indicated sub-MIC concentration of β-peptide **1** (1, 2, or 3 μM) in DPBS. (a) CLSM and corresponding DIC images of PI uptake into *C. albicans* after 45 min incubation in β-peptide solutions of the indicated concentrations. Color represents fluorescence imaging pixel values. Red cells are lysed cells that were excluded from quantitative analysis. (b) Box chart of the distribution of PI uptake in live *C. albicans* as a function of β-peptide concentration and incubation time. Black, red, and blue bars represent 15 min, 30 min, and 45 min incubation times, respectively. The box chart shows median intensity values (solid horizontal line in the box), average intensity (open square in box), 50^th^ percentile values (box outline), 10^th^ and 90^th^ percentile values (whiskers), 1^th^ and 99^th^ percentile values (×), and minimum and maximum intensities (−). Statistical analysis was performed using one-way ANOVA.

### Nuclear and vacuolar membranes are intracellular targets of 14-helical β-peptides

Because β-peptides form pores in the *C. albicans* membrane that mediate molecular translocation into the cytoplasm, it is possible that β-peptides could also cross the cell membrane and act on intracellular targets to promote cell death. To track β-peptide localization inside the cell, we monitored the co-localization of β-peptide **1-Cy5** with organelle-specific dyes. *C. albicans* cells were incubated with organelle-specific dyes, a mixture of β-peptides **1-Cy5** and **1**, and PI. Specifically, FM 4–64 was used to stain the plasma membrane, and SYTO 13 and Mitotracker Green were pre-incubated to stain the nucleus and mitochondria, respectively. MDY-64 was added after removing the β-peptide solution to stain the vacuolar membrane. Fig. 7a shows that β-peptide co-localized with FM 4–64, consistent with prior findings of β-peptide localization to the cell membrane (Walther and Wendland, 2004). Following adhesion to the cell membrane, the β- peptide next translocated into the cytoplasm and associated with the nuclear membrane and the vacuolar membrane. Fig. 7b shows the β-peptide interacting with the nuclear membrane, while Fig. 7c shows β-peptide association with the vacuole. However, β-peptide did not interact with all intracellular membranes. For example, β-peptide did not co-localize with mitochondria (Fig. 7d). These colocalization studies suggest that a mechanism involving lysis of nuclear and/or vacuolar membranes may contribute to the antifungal activity of β-peptides.

**Figure 7.**
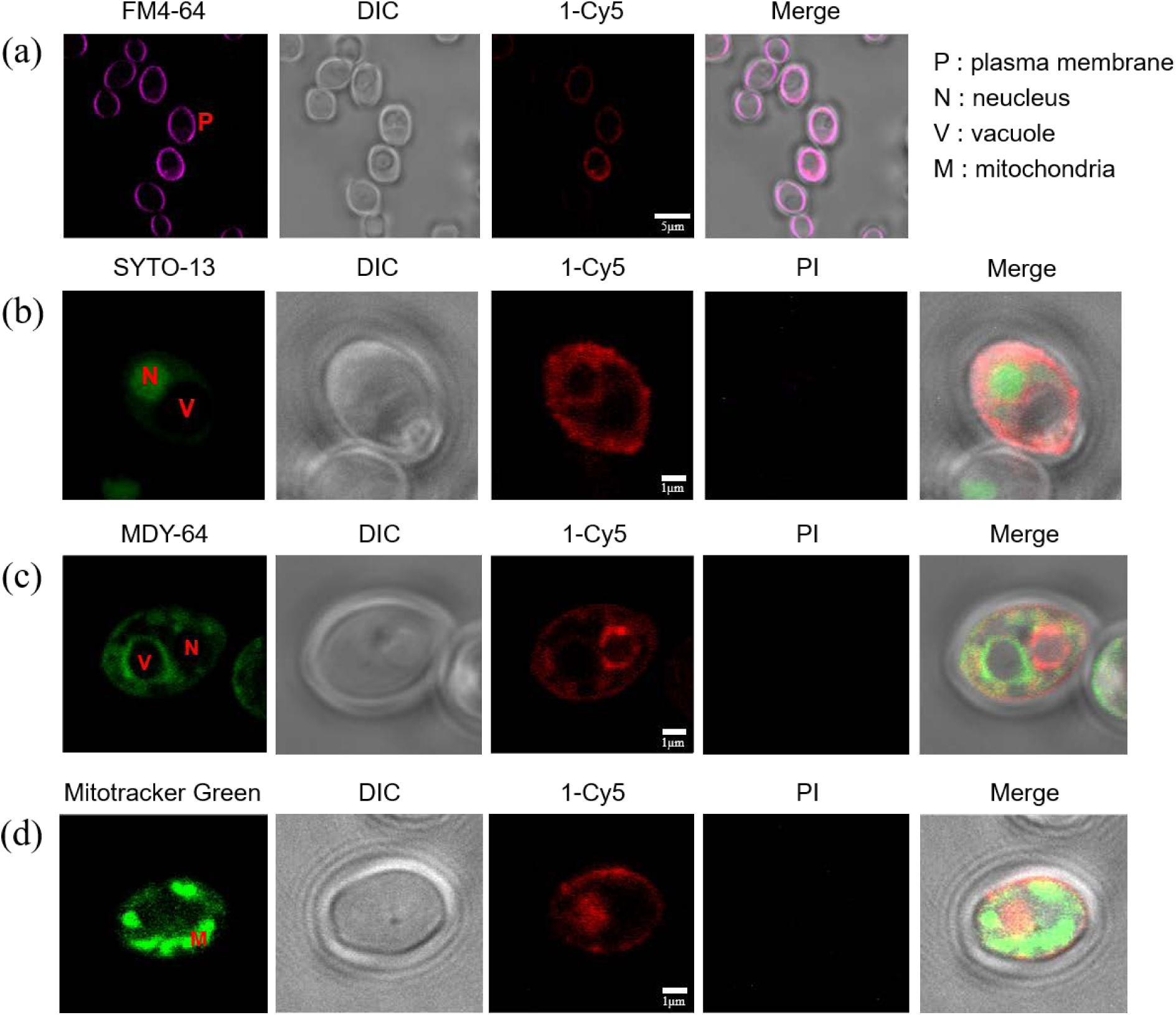
Subcellular localization of β-peptide **1-Cy5**. 2 μM β-peptide **1-Cy5**, 3 μM unlabeled β- peptide **1** and an organelle specific dye (1 μg/mL FM 4–64 for plasma membrane, 2.5 μM SYTO 13 for nucleus, 0.5 μM Mitotracker Green for mitochondria, and 5 μM MDY 64 for intracellular membranes) were incubated with *C. albicans* cells for 5 - 30 min and then imaged. *C. albicans* incubated with β-peptides and (a) FM 4–64, (b) SYTO 13, (c) MDY 64, and (d) Mitotracker green. Scale bar indicates 5 μm for (a) and 1 μm for (b), (c), and (d). P labels the plasma membrane, N the nucleus, V the vacuole, and M the mitochondria.

To determine whether β-peptides also lyse the nucleus and vacuole, we used CLSM to monitor the kinetics of association of β-peptide **1-Cy5** with intracellular organelles. Movie S1 shows that the β-peptides rapidly adhered to the *C. albicans* cell membrane. Next, the cell appeared to shrink, likely as a result of leakage of cytoplasmic contents following poration of the cell membrane. The β-peptide then translocated into the cytoplasm where it adhered to the nuclear and vacuolar membranes. The nuclear membrane was then disrupted, resulting in DNA dispersal throughout the cytoplasm and then the vacuole expanded until the vacuolar membrane disrupted. This sequence of events occurred over 15 min and was observed in all *C. albicans* cells examined. Together, these results establish the plasma membrane, nuclear membrane, and vacuole membrane as targets for amphiphilic 14-helical β-peptides that exhibit antifungal activity.

## DISCUSSION

The mode of action of helical antimicrobial peptides involves electrostatic interactions between a randomly-oriented amphiphilic peptide and the outer membrane of a target cell at the adhesion stage. Upon binding, AMPs adopt a stable and primarily helical secondary structure and self-associate on the outer membrane until a threshold concentration is reached, resulting in cell membrane disruption, either by formation of structured pores or less-organized carpets (Shai, 1999; Yeaman and Yount, 2003). Here, we demonstrate that peptidomimetic 14-helical β-peptides that display antifungal activity lyse the cell membrane by pore formation, allowing the β-peptides to translocate into the cytoplasm where they disrupt the nuclear and vacuolar membranes, resulting in cell death. First, we demonstrated that β-peptide permeabilization of SUVs depends upon SUV composition as well as β-peptide hydrophobicity. Based on prior work showing that β- peptide activity depends on hydrophobicity, charge, and helical stability, we predicted that these interactions would also influence β-peptide permeation of SUVs (Lee et al., 2014; Raman et al., 2015). Indeed, calcein leakage from SUVs correlated with β-peptide hydrophobicity. Neutral charged PC:PE (60:40) SUVs exhibited greater calcein leakage than negatively charged PC:PG (60:40) SUVs after exposure to cationic β-peptides. Hydrophobic β-peptides **1** and **2** induced greater calcein leakage than the more hydrophilic β-peptide 3. Thus, β-peptide hydrophobicity appears have a more significant influence on the ability to disrupt neutral SUVs than anionic SUVs. Hydrophobic β-peptide **2** induced similar calcein leakage from both neutral and negatively charged SUVs at 10 μg/mL peptide concentration, while less hydrophobic β-peptides **1** and **3** resulted in greater calcein leakage from negatively-charged SUVs than neutral SUVs.

The ability of β-peptides to permeabilize SUVs correlated with binding to the cell membrane of live *C. albicans* and to antifungal activity. Wieprecht et al. showed that hydrophobic derivatives of magainin 2a caused greater calcein leakage from POPC/POPG LUVs than native magainin 2a (Wieprecht et al., 1997). Jin et al. demonstrated that the anti-microbial activities of sheet-structured 18-mer and helical 21-mer oligopeptides correlated with calcein leakage from POPC/POPG and POPE/POPG LUVs (Jin et al., 2005). Similarly, our results illustrate that the more active antifungal β-peptides **1** and **2** induce significantly greater calcein leakage from PC:PE(60:40) and PC:PE(80:20) SUVs than the less active β-peptide **3**.

SUV studies also demonstrated cooperative binding of antifungal 14-helical β-peptides in a manner consistent with pore formation. In the rapid accumulation stage, the β-peptides adhere to the cell membrane with a low partition coefficient. After reaching a threshold concentration, the β- peptides insert into the core of lipid bilayer and permeabilize the membrane in a cooperative manner, resulting in a higher partition coefficient (Huang, 2000; Shai, 1999). Rapaport et al. reported a critical concentration of 35 nM for the pore-forming pardaxin derivative (C^1^-NBD-par) against PC SUVs (Rapaport and Shai, 1991). Gazit et al. demonstrated that two helical peptides derived from the pore-forming *Bacillus thuringiensis* serotype israelensis (Bti) toxin exhibited similar critical concentrations and partition coefficients to the full length Bti toxin (Gazit et al., 1994). The NBD-labeled 14-helical β-peptides used in this study exhibited high partition coefficients and low critical concentrations against model membranes (Table 2).

Our live *C. albicans* studies also support a pore-forming mechanism. CLSM measurement of fluorescence accumulation using mixtures of β-peptides **1-Cy5** and **1** showed cooperative binding to the plasma membrane of live *C. albicans* near the MIC of β-peptide **1**. This cooperative adhesion is likely related to a transition from the low to the high partition coefficient observed in the SUVs. Interestingly, β-peptide **1** permitted PI translocation across the cell membrane in live *C. albicans* at concentrations below the MIC. The intracellular concentration of PI depended on the β-peptide concentration and incubation time, but some membrane permeabilization was observed without resulting in cell death. Conversely, β-peptide concentrations above the MIC resulted in rapid entry of high concentrations of PI. Together, these synthetic SUV and live cell studies are consistent with a pore-forming mechanism of β-peptide permeabilization of the *C. albicans* cell membrane.

Mechanisms involving both plasma membrane permeation and intracellular targets have been reported. For example, buforin 2 forms toroidal pores in bacterial cell membranes, then translocates into the cytoplasm and binds to DNA and RNA (Kobayashi et al., 2004; Park et al., 1998; Takeshima et al., 2003). Once inside the cell, 14-helical β-peptides can disrupt the nuclear and vacuole membranes, as determined by time-lapse co-localization of Cy5-labeled β-peptides with organelle-specific fluorescent dyes. After translocating through the cell membrane, the β- peptides bind and permeabilize nuclear and vacuole membranes, sequentially (Movie S1). We show that β-peptide-induced disruption of cellular membranes, including the plasma membrane, nucleus, and vacuole, results in cell death. We predict that the potential of these compounds to disrupt multiple cell membranes could help impede development of resistance mechanisms involving changes in membrane composition.

### Significance

Peptidomimetic AMPs have tremendous potential as antimicrobial agents. However, the mechanisisms of their activity and selectively are not well-understood, limiting efforts to design more effective peptidomimetic AMPs. Our results establish that 14-helical β-peptides disrupt artificial membranes and live *C. albicans* cell membranes. β-Peptides bind to membranes via hydrophobic and electrostatic interactions, and upon reaching a critical threshold concentration insert into and permeabilize the cell membrane in a cooperative manner. Once inside *C. albicans* cells, β-peptides bind and permeabilize nuclear and vacuolar membranes sequentially, leading to cell death. This understanding of the mechanisms of antifungal activity will facilitate design and development of AMP peptidomimetics, including 14-helical β-peptides, for antifungal applications.

## Acknowledgements

The authors thank Sarah Swanson at the UW-Madison Newcomb Imaging Center and Julie Last and the UW-Madison Materials Science Center for imaging assistance. This research was supported by NIH grants RO1AI092225 and R21AI127442. The ESI mass spectrometry for peptide identification was supported by NIH grant 1S10OD020022-1.

## Author Contributions

M.-R.L. and S.P.P. designed the experiments. M.-R.L. and N.R. performed the experiments. M.- R.L., N. R., P.O.-B., D.M.L., and S.P.P. analyzed the data and wrote the manuscript.

## Methods

- Key Resources Table
- Contact for reagent and resource sharing
- Materials
- Methods Details

- Synthesis of Fluorescently-labeled β-peptides
- Measurement of MIC of antifungal peptides
- Preparation of small unilamellar vesicles
- Calcein leakage
- Partition Coefficient measurement
- Adhesion of C. albicans to glass plates for microscopy
- Confocal laser scanning microscopy
- Super resolution structure illumination microscopy
- Characterization of PI uptake by cells
- β-peptide with cellular organells
- Time lapse tracking of β-peptide

### Key Resource Table

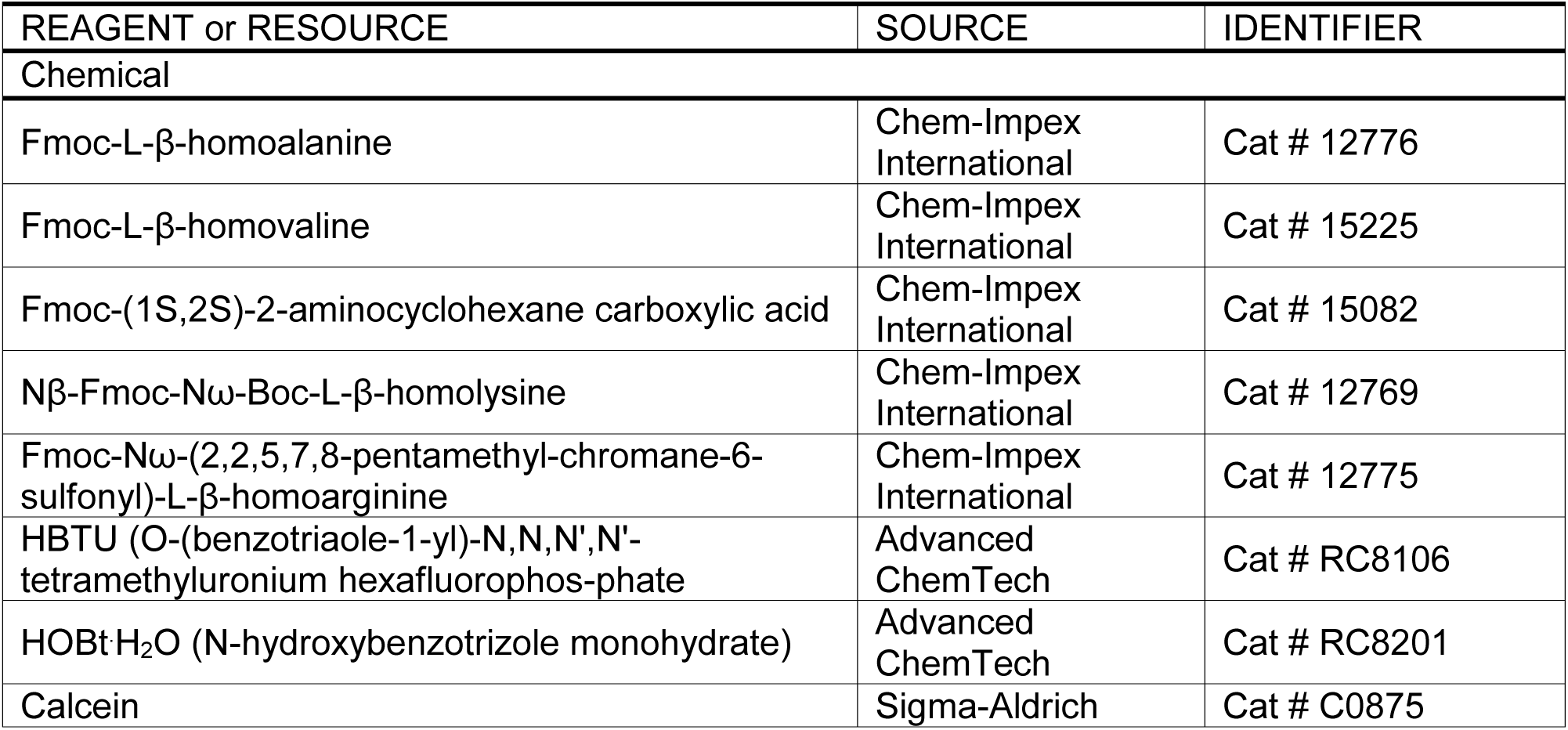
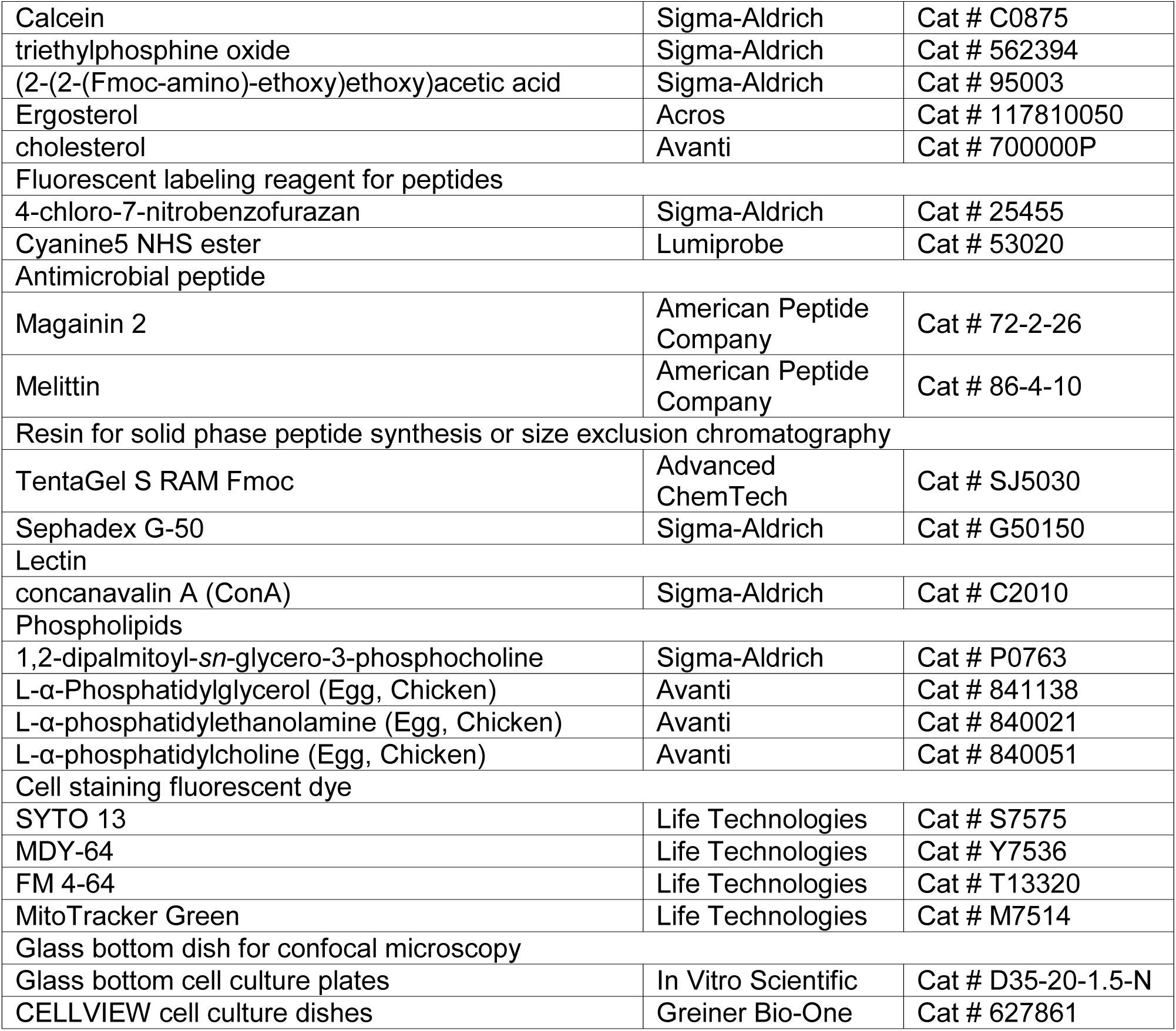

### Contact for Reagent and Resource Sharing

Further information and requests for resources and reagents should be directed to and will be fulfilled by the lead contact, Sean Palecek

### Synthesis of fluorescently-labeled β-peptides

β-peptides were synthesized as previously reported by microwave-assisted protocols of solid-phase peptide synthesis(Lee et al., 2014). NDB and Cy5 labeling were performed after coupling and deprotection of the last β-amino acid. For NBD labeling, NBD-Cl (3 eq), and DIEA (6 eq) were premixed in DMF and then mixed with the β-peptide resin. After stirring overnight at room temperature, the resin was washed with DMF (5 times), CH_2_Cl_2_ (5 times), and DMF (5 times). Prior to Cy5 labeling, β-peptides were coupled with an ethoxy-containing tether, (2-(2-(Fmoc-amino)ethoxy)-ethoxy)acetic acid (3 eq), and then the resin was washed with DMF (5 times), CH_2_Cl_2_ (5 times), and DMF (5 times). Cy5-NHS (2 eq), and DIEA (4 eq) were dissolved in DMF and then mixed with the β-peptide resin. After stirring overnight at room temperature, the resin was washed with DMF (5 times), CH_2_Cl_2_ (5 times) and DMF (5 times). After the NBD and Cy5 labeling, the resin was washed with CH_2_Cl_2_ and then the peptide was cleaved from the resin by addition of TFA containing H_2_O (2.5%) and triisopropylsilane (2.5%) for 1 to 2 h. The crude products of dye-labeled β-peptides were purified by preparative Rβ-HPLC with a gradient of 25–73% CH_3_CN in water containing 0.1% TFA over 35 min. The peptides were validated using mass spectrometry. **1-NBD**: HRMS (m/z, ESI) calcd for C_66_H_112_N_16_O_12_ [M+2H]^2+^ 661.4396, measured 661.4417, **2-NBD**: HRMS (m/z, ESI) calcd for C_66_H_112_N_22_O_12_ [M+TFA+2H]^2^+ 760.4452, measured 760.4482, **3-NBD**: HRMS (m/z, ESI) calcd for C_60_H_100_N_16_O_12_ [M+2H]^2^+ 619.3926, measured 619.3947, 1-Cy5: MALDI-TOF (m/z) calcd for C_98_H_159_N_16_O_13_ [M]+ 1768.2, measured 1768.5.

### Measurement of Minimum Inhibitory Concentration (MIC) of Antifungal Peptides

The MICs of fluorophore-labeled β-peptides (**1-NBD**, **2-NBD**, **3-NBD**, and **1-Cy5**), magainin 2, and melittin against *C. albicans* (SC5314) were determined in accordance with the broth microdilution assay suggested by the Clinical and Laboratory Standards Institute, including quantitative XTT cell metabolic assessment (Karlsson et al., 2009). MIC was calculated as the minimum concentration that resulted in at least 90% decrease in cell viability. Peptides were assayed in RPMI medium containing 1% DMSO.

### Preparation of Small Unilamellar Vesicles

The total phospholipid was maintained at 36 μmol to prepare calcein-encapsulating small unilamellar vesicles (SUVs): PC/PE (21.6/14.4 μmol, 60:40), PC/PE (28.8/7.2 μmol, 80:20), PC/PG (21.6/14.4 μmol, 60:40). The desired amounts of phospholipids were dissolved in chloroform (5 mL) in a round bottom flask and then chloroform was evaporated to form multilamellar layers of phospholipids. After drying the round bottom flask under vacuum for 1 h, 70 mM calcein in TBS buffer (10 mM Tris·HCl, 150 mM NaCl, 0.1 mM EDTA, pH 7.4, 4 mL) and 10 N NaOH (3 eq) was added into the flask and then the solution was vortexed to form a suspension of multilamellar vesicles. This suspended solution was sonicated for 30 to 45 min in an ice bath using an ultrasonicator (550 Sonic Dismembrator, Fisher Scientific). Unilamellar vesicles were purified by Sephadex G-50 size exclusion chromatography (L/D = 26) under gravity. After purification, the vesicle composition was measured by ^31^P NMR and the effective vesicle size was quantified by dynamic light scattering (DLS). The DLS of a 1 mL purified SUV solution was measured at 25 ^o^C in a glass cell in a decahydronaphthalene environment. Light scattering was measured at a 90° angle using a 100 mW, 532 nm laser (Coherent Compass 315M-100) and BI-9000AT digital autocorrelator (Brookhaven Laser Light Scattering System, Brookhaven Instruments Corporation). The effective vesicle size was measured between 50 - 90 nm (Table S1). To measure β-peptide partition coefficients, unilamellar vesicles were prepared as described for calcein-encapsulating vesicles. PC/PE (60:40) and PC/PG (60:40) vesicles were prepared at a 36 μmol scale in TBS buffer (4 mL). PC/PE/ergosterol (22.5/13.5/9 μmol, 50:30:20) vesicles were prepared at a 45 μmol scale in TBS buffer (9 mL). After purification, the vesicle composition and phospholipid concentration were measured by ^31^P NMR and ^1^H NMR and the effective size of vesicles was measured by DLS. The effective vesicle size was measured between 40 - 80 nm (Table S2).

### Calcein leakage

The desired concentrations of β-peptides (**1** - **3**), magainin 2, and melittin in TBS buffer were added to a 96 well plate (75 μL) with positive (2% Triton X100 in TBS buffer) and negative (TBS buffer) lysis controls. SUVs (75 μL, 260 μM) were added into the 96 wells of the plate and the fluorescence intensity was recorded (ext. 495 / em. 515 nm) over 1 h at 80 s intervals using a FL800 Universal Microplate Reader) (Bio-Tek Instruments). The calcein leakage (%) was calculated as

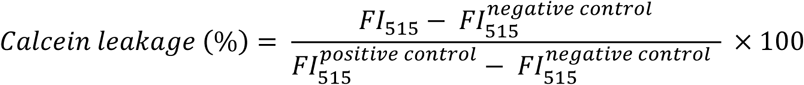

where *FI*_515_, 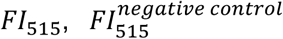 and 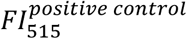 are the average fluorescence intensities at 515 nm of SUVs treated with peptides, TBS, and Triton X100, respectively. Duplicate samples were analyzed for experiments repeated on at least three different days.

### Partition coefficient measurement

All NBD labeled β-peptide stock solutions were prepared in DMSO (20 μM). SUV solution (792 μL) and peptide stock solution (8 μL) were mixed in a fluorescence cuvette and then reacted for 2 min before the recording fluorescence (ext. 468) spectrum from 480 - 600 nm (F-4500 Fluorescence Spectrometer, Hitachi). Experiments were performed in triplicate using freshly-prepared SUVs on different days. After recording fluorescence spectra, the partition coefficients were calculated as reported previously (Pouny et al., 1992; Rapaport and Shai, 1991). Briefly, the binding isotherms were assessed as partition equilibria using the following equation:

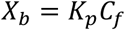

where *X_b_, K_p_* and *C_f_* are the molar ratio of bound peptide per total lipid, the partition coefficient, and the equilibrium concentration of free peptide in the solution, respectively. In order to calculate *X_b_, F*_∞_ (the estimated maximum fluorescence intensity when vesicles are saturated by peptide) was calculated from the peptide titration curve. The *f_b_* (fraction of bound peptide) was calculated from the following equation:

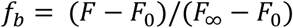

where *F*, and *F*_0_ are the fluorescence intensity of bound and unbound NBD-peptide, respectively. *X_b_* equations were corrected by the fraction of phospholipid accessible to the 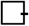 peptide (60% of total lipid). Thus, the binding isotherm was represented as:

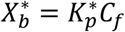

where 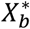 and 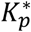 are the molar ratio of bound peptide per 60% of total lipid and the estimated surface partition constant, respectively.

### Adhesion of *C. albicans* to glass plates for microscopy

A single colony of *C. albicans* (SC5314) was suspended in 5 mL YPD medium and grown overnight at 32.5 ^o^C. The cells were collected by centrifugation (2000 rpm, 5 min) and washed with DPBS buffer. ConA (2 mg/mL in DPBS, 500 μL) was adsorbed to the glass bottomed cell culture dish for 24 h at 4 ^o^C. After incubation, the dish was rinsed with DPBS. The *C. albicans* cells (10^6^ - 10^7^ cells/mL in 400 μL) were seeded on ConA-coated plates and allowed to attach for 30 min prior to CLSM and SR-SIM microscopy.

### Confocal laser scanning microscopy

*C. albicans* cells adhered to a glass-bottomed dish were incubated with **1-Cy5** (0.5 μM) and varying concentrations of β-peptide **1** (0, 0.5, 1.5, 3.5, 6.5 μM) in DPBS (300 μL). After 3 min incubation, cells were imaged by CLSM (Zeiss LSM 510) with a 100x objective. The excitation and emission filter were selected, excitation filter: 633 nm and emission filter: BP 650–710 IR, respectively. Images were processed using Zeiss LSM image browser.

### Super resolution structure illumination microscopy (SR-SIM)

*C. albicans* cells adhered to a glass-bottomed dish were incubated with 2 μM **1-Cy5** in DPBS (300 μL) for 3 min and then cells were imaged by CLSM (Zeiss LSM 780) with an alpha planapochromat 100x/1.46 oil objective equipped with a 642 nm laser. Immersion oil was applied on the objective to minimize refractive index mismatch and spherical aberration. 3D SR-SIM image stacks of 3.84 μm thickness were acquired with 0.133 μm z-steps and 15 images (5 rotations per angle) per z-section. Images were processed using Zeiss ZEN Software.

### Characterization of PI uptake by cells

10 μg/mL PI and β-peptide 1 (1, 2, or 3 μM) in DPBS (300 μL) were added to cells adhered to a glass-bottomed dish. Images were acquired for 45 min with 3 min intervals by CLSM (Zeiss LSM 780) with a 100x objective. The excitation and emission filter were selected, excitation filter: 561 nm and emission filter: 575–600 nm bandpass filter, respectively. Images were processed using Zeiss ZEN software.

### Characterization of co-localization of β-peptide with cellular organelles

*C. albicans* cells were adhered to a glass-bottomed dish as described above. **For FM 4-64/1-Cy5:** A solution of 1 μg/mL FM 4-64, 2.0 μM β-peptide **1-Cy5**, and 3.0 μM β-peptide **1** in DPBS (300 μL) were added to the cells and incubated for 5 min. Then, the confocal images were acquired. **For SYTO 13/1-Cy5:** A solution of 2.5 μM SYTO 13 in DPBS (300 μL) was added to the cells and incubated for 20 min. After incubation, the SYTO 13 solution was removed from the dish and then a solution of 2.5 μM SYTO 13, 2.0 μM β-peptide **1-Cy5**, 3.0 μM β-peptide **1**, and 1 μg/mL PI in DPBS (300 μL) was added to the dish. After 5 - 10 min, confocal images were acquired. **For MDY-64/1-Cy5:** A solution of 2.0 μM β-peptide **1-Cy5** and 3.0 μM β-peptide **1** in DPBS (300 μL) was added to the cells and incubated for 10 min. After incubation, the peptide solution was removed from the dish and the cells were washed 2 times with DPBS. A solution of 5 μM MDY-64 and 1 μg/mL PI in 10 mM HEPES containing 5% glucose, pH 7.4 (300 μL) was added to the dish. Immediately after, confocal images were acquired. **For MitoTracker Green/1-Cy5**: A solution of 0.5 μM MitoTracker in DPBS (SOO μL) was added to the cells and incubated for 30 min. After incubation, the MitoTracker solution was removed from the dish and then a solution of 2.0 μM β- peptide **1-Cy5**, 2.0 μM β-peptide **1**, and 1 μg/mL PI in DPBS (300 μL) was added to the dish. After 5 - 10 min, confocal images were acquired. Images were acquired by CLSM (Zeiss LSM 510) using a 100x objective with Sx zoom. The excitation wavelengths and emission filters were selected as followed: SYTO 1S (ext. 488 nm, filter: BP 500-530 IR), Cy-5 (ext. 633 nm, filter: BP 650-710 IR), FM 4-64 and PI (ext. 561 nm, filter: BP 575-615 IR), MDY-64 and MitoTracker Green (ext. 458 nm, filter: BP 480-520 IR). Images were processed using Zeiss LSM image browser.

### Time lapse tracking of β-peptide

A solution of 1 μg/mL Hoechst in DPBS (300 μL) was added to cells adhered to a glass-bottomed dish and incubated for 30 min. After incubation, the Hoechst solution was replaced with the solution of 1 μg/mL Hoechst and 0.5 μM Mitotracker green in DPBS (300 μL) and incubated for an additional 20 min. Then a solution of 1 μg/mL PI, 2.0 μM β-peptide **1-Cy5**, and 3.0 μM β-peptide **1** in DPBS (300 μL) was added to the dish. Confocal images were acquired for Hoechst, Mitotracker green, PI and Cy5 every 40 s. Cells were imaged using CLSM (LSM 700) with a 60x objective. The excitation wavelength and emission wavelength filters were selected as followed: Hoechst (ext. 405 nm, filter EM 500–530 nm), Cy5 (ext. 639 nm, filter EM 650–710 nm), Mitotracker green (ext. 488 nm, filter EM 480–520 nm), and PI (ext. 561 nm, filter EM 575–615 nm). Images were processed using Zeiss ZEN software.

